# Stimulus representations in visual cortex shaped by spatial attention and microsaccades

**DOI:** 10.1101/2023.02.25.529300

**Authors:** Karthik Srinivasan, Eric Lowet, Bruno Gomes, Robert Desimone

## Abstract

Microsaccades (MSs) are commonly associated with spatially directed attention, but how they affect visual processing is still not clear. We studied MSs in a task in which the animal was randomly cued to attend to a target stimulus and ignore distractors, and it was rewarded for detecting a color change in the target. We found that the enhancement of firing rates normally found with attention to a cued stimulus was delayed until the first MS directed towards that stimulus. Once that MS occurred, attention to the target was engaged and there were persistent effects of attention on firing rates for the remainder of the trial. These effects were found in the superficial and deep layers of V4 as well as the lateral pulvinar and IT cortex. Although the tuning curves of V4 cells do not change depending on the locus of spatial attention, we found pronounced effects of MS direction on stimulus representations that persisted for the length of the trial in V4. In intervals following a MS towards the target in the RF, stimulus decoding from population activity was substantially better than in intervals following a MS away from the target. Likewise, turning curves of cells were substantially sharper following a MS towards the target in the RF. This sharpening appeared to result from both a “refreshing” of the initial transient sensory response to stimulus onset, and a magnification of the effects of attention in this condition. MSs to the target also enhanced the neuronal response to the behaviorally relevant target color change and led to faster reaction times. These results thus reveal a major link between spatial attention, object processing and its coordination with eye movements.

## INTRODUCTION

In the standard covert attention paradigm (Posner, 1980) subjects hold fixation on a central spot while directing attention to an extrafoveal stimulus, ignoring distractors. This paradigm has been a mainstay of studies in spatial attention, because it separates the physiological role of brain systems involved in the control of eye movements from those that either control attention or are modulated by attention (Desimone and Duncan, 1995; Maunsell, 2015; Moore and Zirnsak, 2017). Yet, studies of covert attention in humans and monkeys have shown that, in spite of instructions to maintain fixation, subjects often make small eye movements towards and away from the attended stimulus, and these fixational eye movements or microsaccades, are often correlated with performance on the subject’s task (Denison et al., 2019; Engbert, 2006; Engbert and Kliegl, 2003; Hafed et al., 2015; Hafed, 2013; Hafed and Krauzlis, 2012; Intoy et al., 2021; Laubrock et al., 2005; Martinez-Conde et al., 2013, 2009; Poletti et al., 2017; Poletti and Rucci, 2013; Tian et al., 2016; Veale et al., 2017; Willeke et al., 2019) as well as the gain of neuronal responses (Bair and O’Keefe, 1998; Chen et al., 2015; Hafed, 2013; Herrington et al., 2009; Snodderly et al., 2001). Thus, in spite of the widely held view that the neural bases for covert attention and saccades are separable, these studies suggested that there may be, in fact, a strong relationship between the two.

We previously reported (Lowet et al., 2018) that in monkeys tested in a covert attention paradigm, neurons in area V4 and IT cortex had enhanced responses during attention to a stimulus in the receptive field (RF), but this enhancement was restricted to the time period immediately following a microsaccade (MS) towards the cued stimulus. No such enhancement was found following a MS back to the fixation target. This strong relationship between MS and the attentional modulation of responses was missed in our prior studies because we had only looked for relationships to MS in general, and not specifically to MS towards versus away from the attended stimulus. However, other studies have reported that attentional modulation of neural responses in primates can occur even during intervals in a covert attention task with no MS. One caveat is that MS could be missed if the eye movement tracking signal is noisy or has poor resolution, and this could give the appearance that neuronal effects can occur without MS. However, one recent study (Yu et al., 2021) of the superior colliculus (SC) with high resolution eye tracking confirmed that the enhancement of responses by attention was strongly modulated by MS towards or away from the attended stimulus, but it also found that attentional enhancement of firing rates could occur even during intervals without MS.

Another recent covert attention study in humans with eye-tracking using EEG showed that the lateralization of alpha power co-varied with microsaccades (Liu et al., 2021), with lateralization being stronger in trials with MS towards an attended stimulus held in working memory. However, similar to the (Yu et al., 2021) study, they too report that such modulations of alpha occur even in trials without MS, thus concluding that MS modulate alpha, but are not obligatory for the modulation of alpha by attention. Are MS required for the attentional enhancement of neuronal responses or not? One limitation of these studies is that they did not examine the interval immediately preceding the MS-free interval, and we previously reported that task intervals that appear to be free of MS were often immediately preceded by a MS, which could mask a relationship between firing rates and MS if one only examined intervals that were putatively MS-free. Thus, the question of whether the modulation of neuronal responses by attention requires a MS towards the attended stimulus remains open.

To help resolve this question, in the present study we have undertaken a new analysis of neuronal responses in areas V4, IT cortex, and the ventrolateral division of the pulvinar during a covert attention task. Each trial in the task could last up to several seconds, and thus contained several intervals with MS. The stimulus was either preceded by or followed by an attentional cue. We found that following the very first MS after either stimulus onset or the attentional cue, firing rates were enhanced by attention almost exclusively according to whether the MS was directed towards or away from the attended stimulus, consistent with what we found previously. On trials where the first MS was made away from the cued stimulus in the RF, attentional effects were delayed until the second MS, which was directed to the cued stimulus. In all MS triggered intervals following the first MS, there was a sustained enhancement of firing rates if attention to the stimulus in the RF was maintained, although the enhancement was greater for MS directed towards the target. This finding suggests that the first MS towards an attended stimulus can initiate a period of sustained attentional enhancement of firing rates. If the first MS after an attentional cue is directed away from the attended stimulus in the RF, the sustained effects of attention will be delayed to the next MS directed towards the cued stimulus interval.

In contrast to these temporally-variable effects of MS on the enhancement of firing rates, we found that stimulus decoding was better and tuning curves were sharper for MS made towards the attended stimulus throughout the entire trials. These results thus reveal a major link between spatial attention, object processing and its coordination with eye movements.

## RESULTS

We used data from the same task described in (Lowet et al., 2018). Briefly, we measured multiunit activity in two awake monkeys (*macaca mulatta*) recorded from laminar microelectrodes inserted in cortical areas V4, IT, and the ventrolateral division of the pulvinar during a spatial attention task (Figure 1A). Monkeys maintained fixation on a central spot while a spatial cue directed attention to one of three extra-foveal stimuli. In some sessions, the cue could occur 500– 700ms after the stimulus (stim-first sessions; Figure 1 top), and in other sessions the cue occurred after the stimulus (cue-first sessions; Figure 1 bottom). After 1,200–1,700ms, the cued stimulus briefly (50ms) changed color, and the monkeys were rewarded for making a saccade to the cued stimulus location. If not specified otherwise, data from stim-first sessions and cue-first sessions were combined. Stimuli were grayscale objects, all in the contralateral hemifield. Eye position was measured with an infrared eye-tracking system and MSs were computed using the algorithm suggested in (Engbert and Kliegl, 2003). MSs occurred at a median rate of 3.29 ± 0.075 Hz and showed two predominant directions for each target location, roughly in opposite directions. About *55%* of the first MS after the cue was made in the direction of the stimulus that was attended and the other *45%* were MS made towards the fixation spot. Subsequent MS were in the opposite direction, followed by a back and forth of MSs between the fixation point and the target.

**Figure 1.**
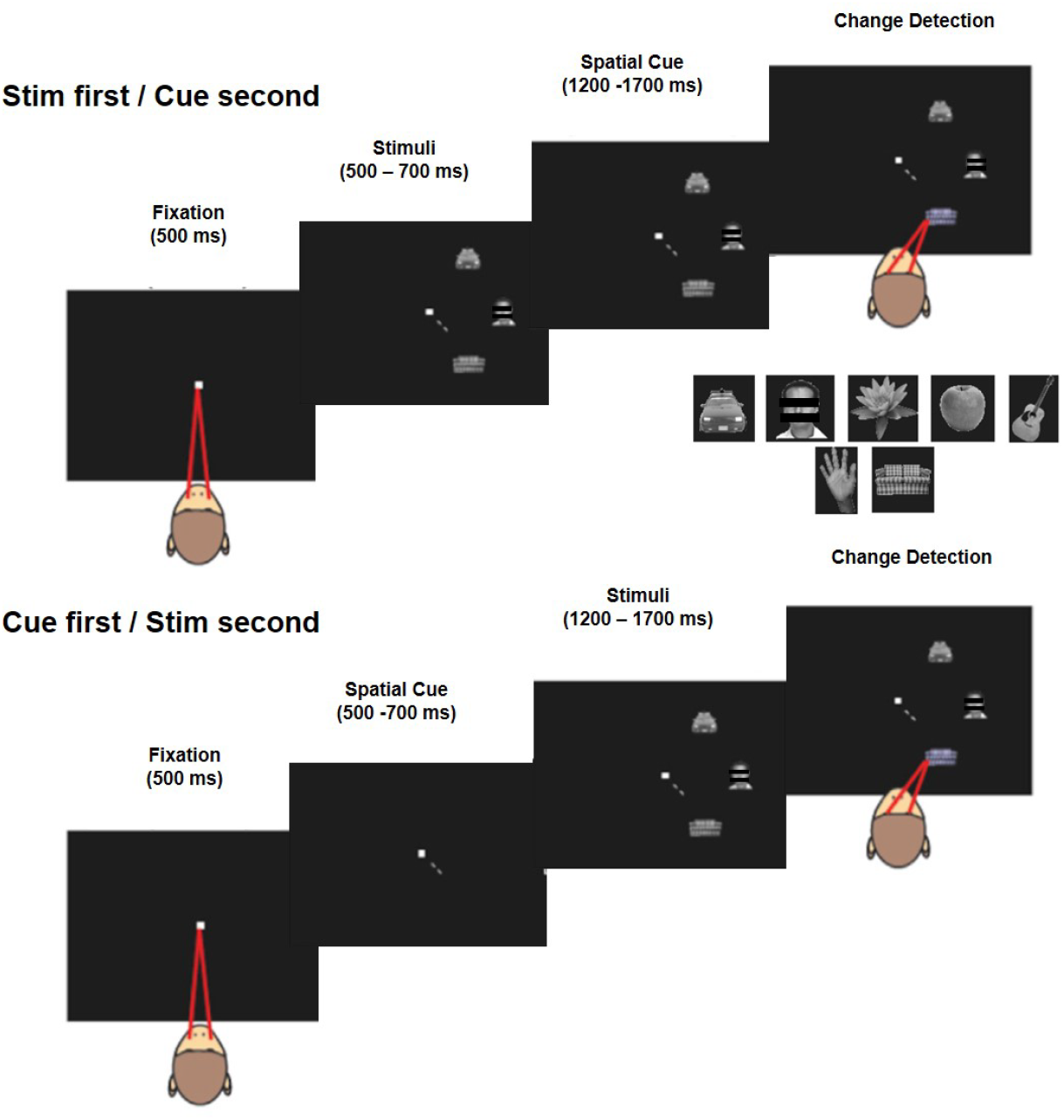
(Top) Task in the stimulus first condition. A central fixation spot first appears followed by a three-stimulus visual array (target and two distractors) in the contralateral hemifield of the recording sites. This is followed by a spatial cue pointing to the target stimulus location that has to be attended. The monkey has to covertly attend to the target while maintaining fixation, followed by a saccade to the target when it changes color. The animal gets rewarded for a saccade to the target and the trial ends. (Bottom) In the cue first condition, the spatial cue appears before the stimulus array appears. (Inset) The set of seven objects that were used in the stimulus array.

To test whether the effects of MSs on the attentional modulation of V4 responses was dependent on the order of MS, we computed firing rates in the 300ms interval following the first MS, contingent on whether the MS was made towards or away from the target. Figures 2A and B shows the population average firing rates for those two conditions, and Figures 2C and D show distribution histograms for all cells of an attentional index for the corresponding MS direction. The average firing rates and the attentional modulation indices (AMI) both show strong enhancement of responses when attention was directed to the RF stimulus, but only when the first MS was directed towards the cued stimulus (*t-test, n=293, p=1*.*05*^*-22*^, *median AMI = 0*.*13±0*.*012)*, consistent with our previous results (Lowet et al., 2018). When the first MS was directed away from the cued stimulus, there was no significant net enhancement of firing rates depending on whether the RF stimulus was attended (*t-test, n=293, p=0*.*649, median AMI = 0*.*004±0*.*01*).

**Figure 2.**
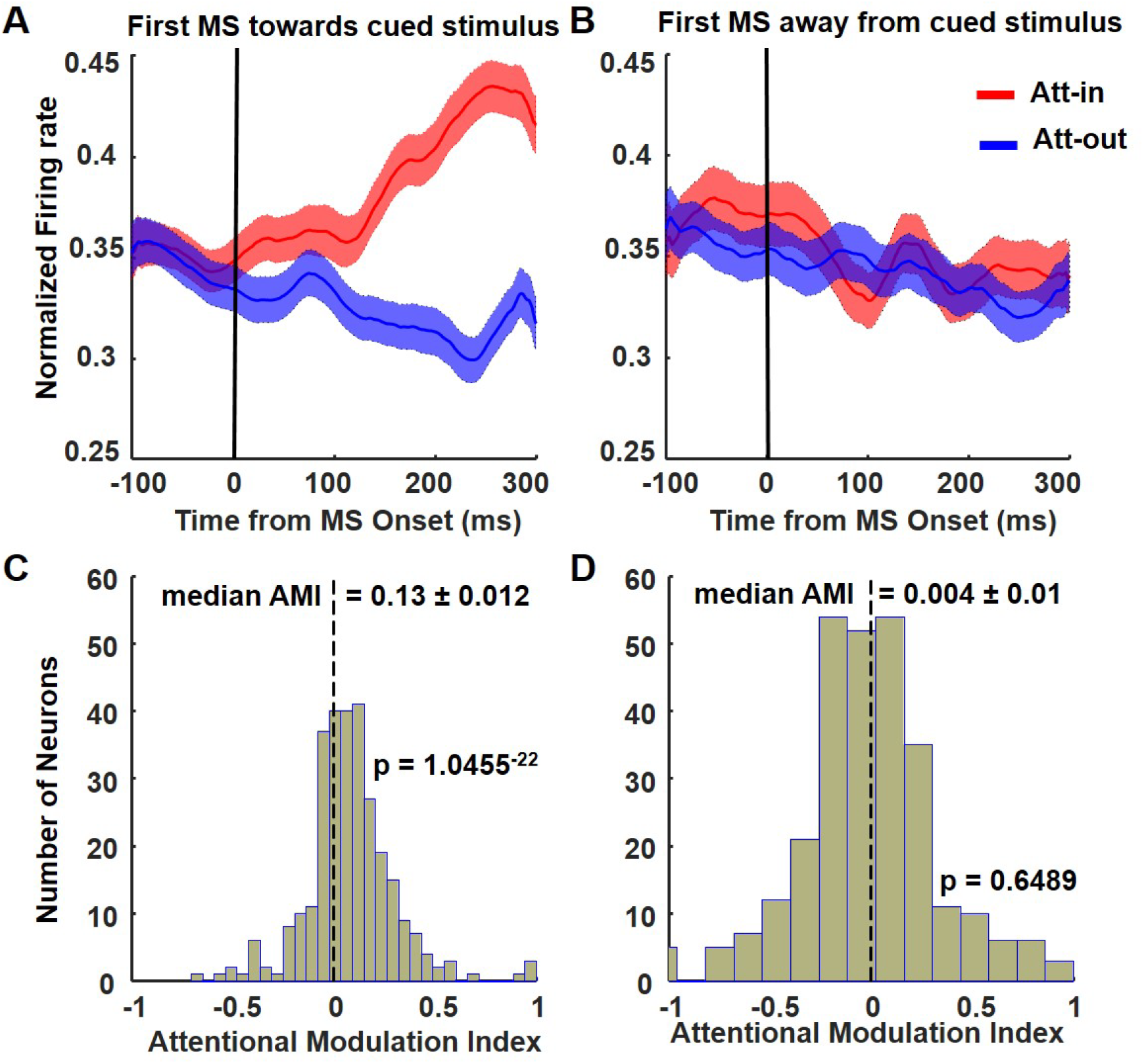
The role of the first MS, and subsequent MSs during the attention period in area V4. (A and B) Normalized population firing rates *(n=293)* combined across monkeys and sessions (*k=34*) locked to the onset of the first MS towards (A) or away from the target location (B) for both Att-in and Att-out conditions, respectively. Only with the first MS towards the RF, do we observe the process of attentional enhancement. (C and D). Histogram distribution of the Attentional Modulation Index for the 300ms following the first MS directed towards (C) or away from the cued stimulus.

One possible explanation for why attentional effects in the first MS interval were only found when that MS was directed towards the cued stimulus in the RF is that attentional effects and/or the deployment of spatial attention to the cued stimulus were delayed until the first MS towards the cued stimulus. If so, then following a first MS away from the cued stimulus in the RF (MS back to fixation), responses should be enhanced on the next MS towards the cued stimulus. Indeed, we found this enhancement on the second MS if the first MS was directed away from the cued stimulus (*t-test, n=293, p=3*.*64*^*-2*^, *median AMI = 0*.*11±0*.*036*) (Fig 3B). If the first MS was directed towards the cued stimulus, then the effects of attention continued, at a lower level, during the next MS directed away from the cued stimulus (*t-test, n=293, p=1*.*97*^*-9*^, *median AMI = 0*.*083±0*.*02*) (Fig 3A). Thus, it appears that the effects of attention on responses are locked to the time when a MS is first made to the cued stimulus, even if it is the second MS after stimulus onset. That MS towards the cued stimulus might be a marker of the first deployment of attention to the target or it might play a mechanistic role in initiating the effects of attention on responses.

**Figure 3.**
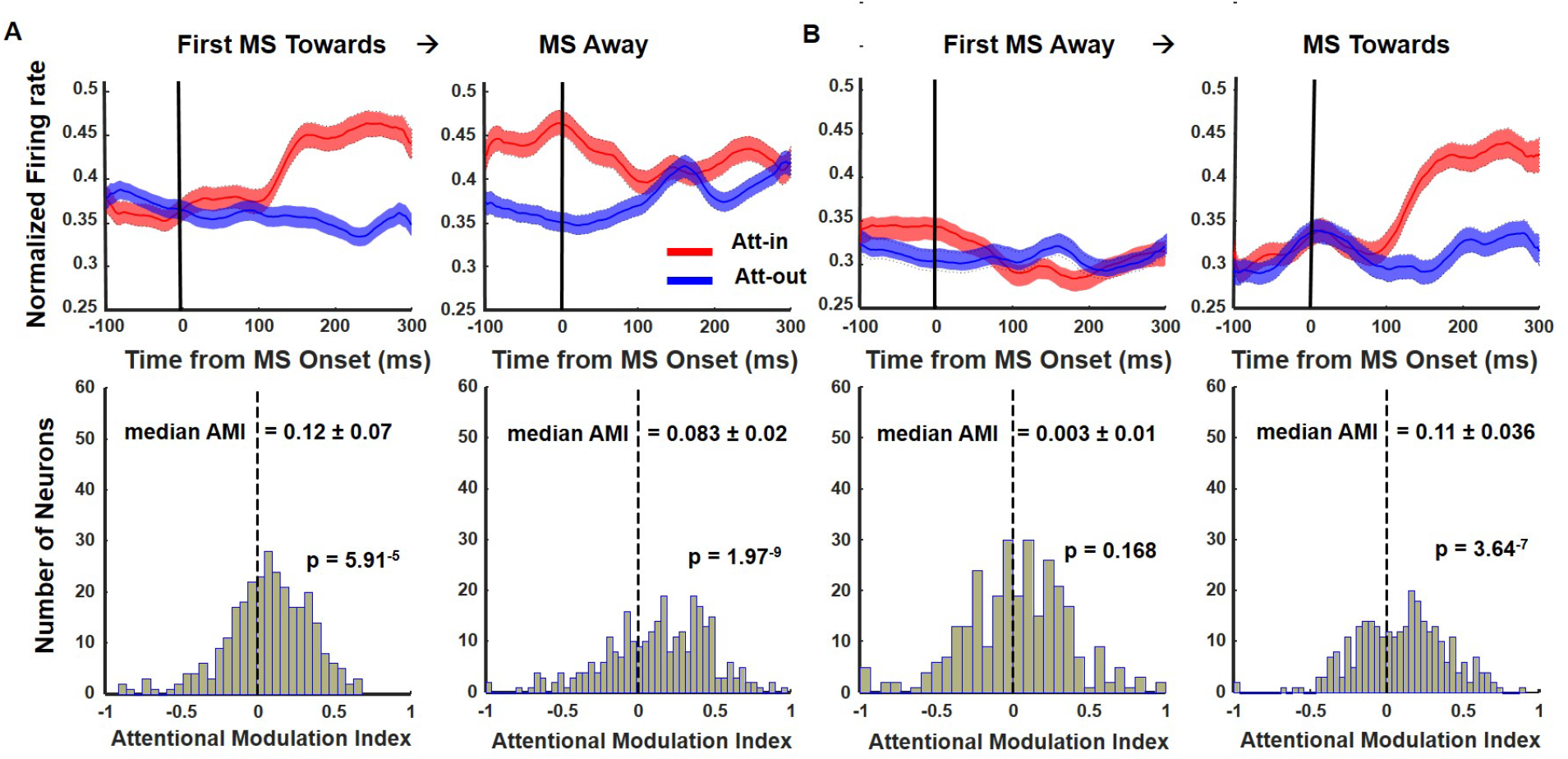
The effects of MS direction on the enhancement of V4 responses with attention depend on MS order. A) If the first MS is directed towards the target stimulus in the RF, responses are enhanced by attention on the first and second MS. Top shows normalized firing rates across the population and bottom show distribution of AMI values. B) If the first MS is directed away from the target stimulus in RF, responses are not enhanced by attention until the next MS towards the target. Top shows normalized firing rates across the population and bottom show distribution of AMI values.

We then examined all subsequent MS that followed the first MS until the end of the trial. Because the target color change could occur at any time from 1,200 to 1,700ms after the cue, there was a variable number of MS intervals after the first MS. To maximize the amount of data for the subsequent MS, we averaged over all of these MS intervals, excluding the interval after the first MS. Figures 4A and C show the population average firing rates and distribution of AMI, respectively, when subsequent MSs were directed towards the cued stimulus (*t-test, n=293, p=3*.*58*^*-21*^, *median AMI = 0*.*14±0*.*015)*. Figure 4B and D show the population average firing rates and distribution of AMI, respectively, when the subsequent MS was directed away from the cued stimulus (*t-test, n=293, p=1*.*25*^*-11*^, *median AMI = 0*.*09±0*.*015)*. For both the MS toward and MS away condition, the firing rates and AMI values were higher when attention was directed into the RF (see Figures for AMI values and statistics), although the AMI was significantly larger following MS directed towards the cued stimulus versus away from that stimulus. Thus, in contrast to the results for the first MS, there was an overall enhancement of firing rates by attention for both directions of subsequent MS. The same pattern of results was found if we separately considered trials when the stimulus appeared first (*total sessions, k =18, number of neurons, n = 161*) and the attentional cue appeared second (Figure S1), or when the cue appeared first (*total sessions, k =16, number of neurons, n = 132*) and the stimulus appeared second (Figure S2).

**Figure 4.**
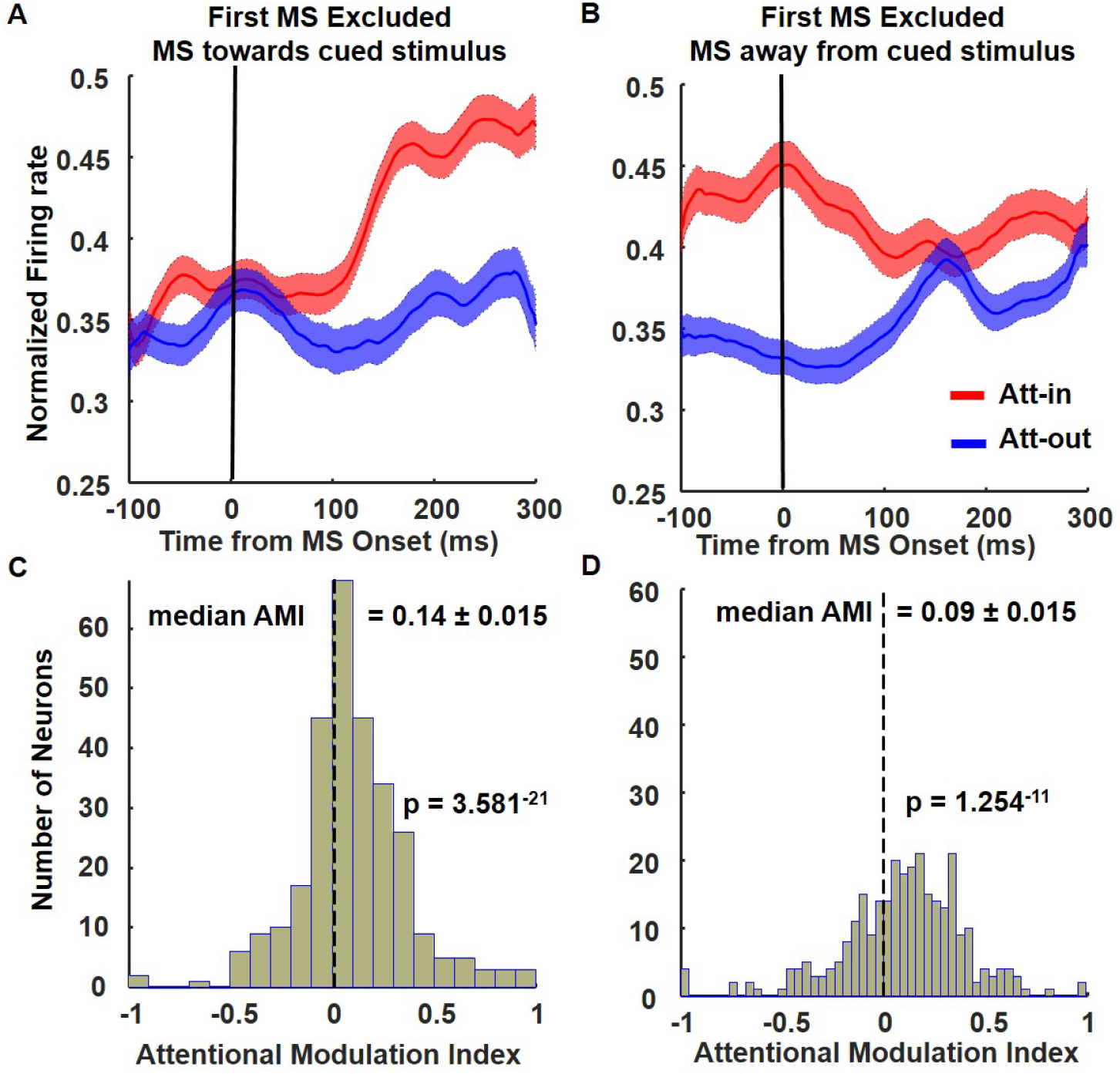
(A and B) normalized population firing rates in V4 (n=293) are locked to all the subsequent MSs excluding the first MS. (C and D) Histograms of AMI values for conditions A and B, respectively.

Although the attentional enhancement of response was maintained under both MS directions after the first MS, the direction of MS appeared to influence the time course of the population firing rate and the magnitude of the attentional enhancement, as shown in the population histograms of Figures 2A and B. When the MS was directed towards the cued stimulus, the effects of attention on firing rates began starting about 100ms after the start of the MS. By contrast, when the MS was directed away from the cued stimulus, the firing rate for the cued stimulus was already elevated compared to the unattended stimulus at the start of the MS, likely because the previous MS was directed towards the attended stimulus. This attentional enhancement slowly diminished over the next 300ms. This slow-to-diminish attentional enhancement following the MS, coupled with the short intervals between MS, may explain why enhancement continues to be positive during MS-away interval even after the first MS.

A similar pattern of attentional enhancement after MS onset was found in both supragranular and infragranular cells. The population firing rate histograms (*n=293*) for cells recorded in both superficial and deep layers of area V4 are shown in Figure S3. The supragranular (*n=109*), input (or granular) (*n=21*), and deep (infragranular) layers (*n=163*) for each session (*k=34 sessions*) were determined by computing the CSD (current source density) at stimulus onset across the 16 channels of the laminar electrode to determine the sources and sinks. The channels with a distinct sink immediately after stimulus onset were considered to be the input/granular layers, and the layers above, and below the input layers as supragranular and infragranular respectively.

On the first MS after the cue, attentional enhancement was limited to the MS-towards condition (Figure S3B supragranular: *t-test, n=109, p=9*.*9*^*-9*^, *median AMI = 0*.*14±0*.*022; infragranular*: *t-test, n=163, p=1*.*62*^*-12*^, *median AMI = 0*.*11±0*.*02)*. However, following the first MS, there was a persistent attentional enhancement for both MS-towards (Figure S3C supragranular: *t-test, n=109, p=1*.*07*^*-10*^, *median AMI = 0*.*17±0*.*025; infragranular*: *t-test, n=163, p=2*.*6*^*-10*^, *median AMI = 0*.*12±0*.*02)*, and MS-away conditions (Figure S3C supragranular: *t-test, n=109, p=2*.*5*^*-6*^, *median AMI = 0*.*13±0*.*024; infragranular t-test, n=163, p=1*.*9*^*-5*^, *median AMI = 0*.*07±0*.*021)* (Figure S3).

In Figure 5, we summarize the AMI values for the MS triggered intervals in the different layers. For both the first MS (Figure 5A), and in intervals with the first MS excluded (Figure 5B), there was significant attentional enhancement for the MS towards condition. We also observed more attentional enhancement in neurons in the superficial (light gray bars) layers than those in the deep layers (red bars).

**Figure 5.**
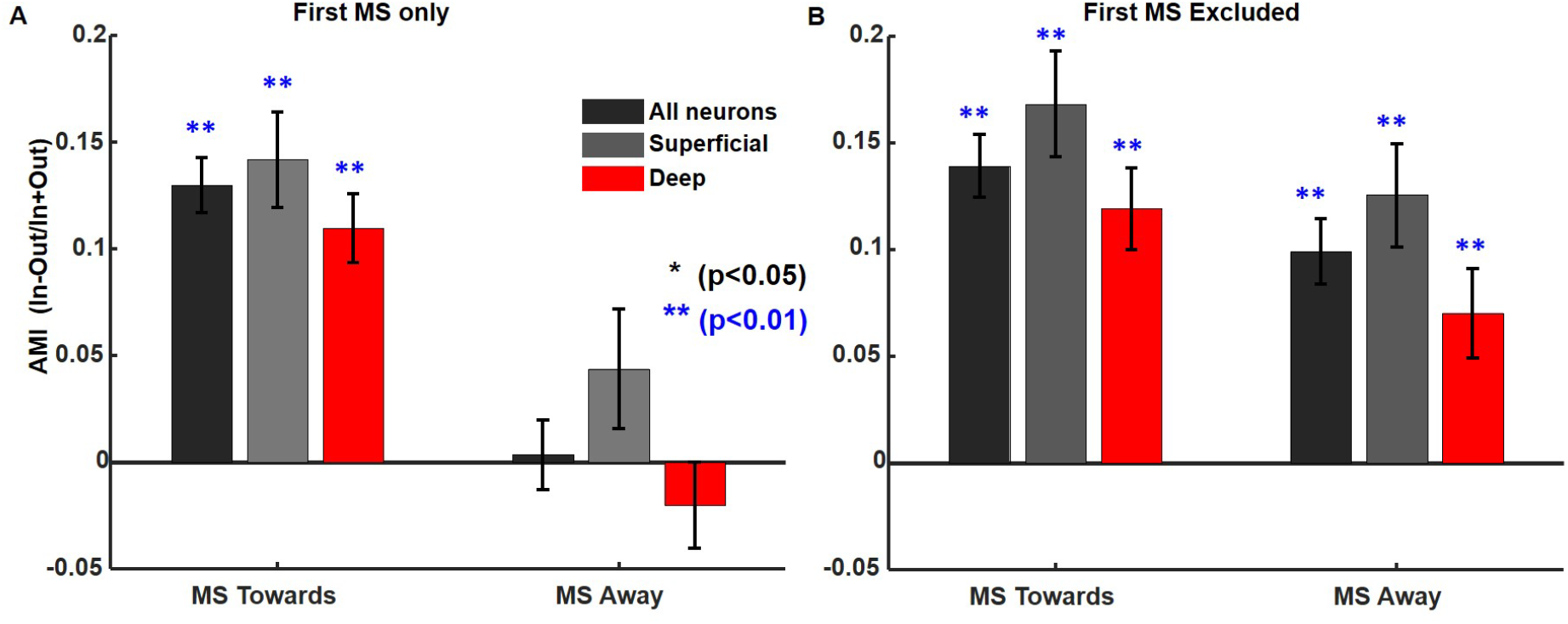
AMI values in all layers combined versus superficial and deep layers separately. A) First MS only. In all neurons, as well as in superficial layers (supragranular) and deep layers (infragranular) there is significant AMI for the MS towards intervals. There is a greater AMI in the superficial layers than in the deep layers. B) Same as in B but with the first MS excluded.

As described above, firing rates in MS intervals subsequent to the first MS were enhanced by attention, but they were nonetheless larger in the MS-towards condition compared to the MS-away condition. We also found that firing rates to the color change of the attended stimulus at the end of the trial were higher for MS-towards intervals (*t-test, n=293, p=1*.*82*^*-21*^, *median AMI = 0*.*1±0*.*01*) (Figure 6A and C) compared to MS-away (*t-test, n=293, p=0*.*19, median AMI = 0*.*008±0*.*01*) (Figure 6B and D). The summary of AMI during the last MS is presented in Figure 6E. This increase in the neural response to the color change was accompanied by faster reaction times (p=0.0056) if the color change occurred during a MS-towards (*RT = 200 ± 52*.*7 ms)* interval compared to MS-away (*RT = 220 ± 56*.*45 ms)* (Figure 6F). The histogram distribution of the RTs and their empirical cumulative distribution are presented in Figure S4.

**Figure 6.**
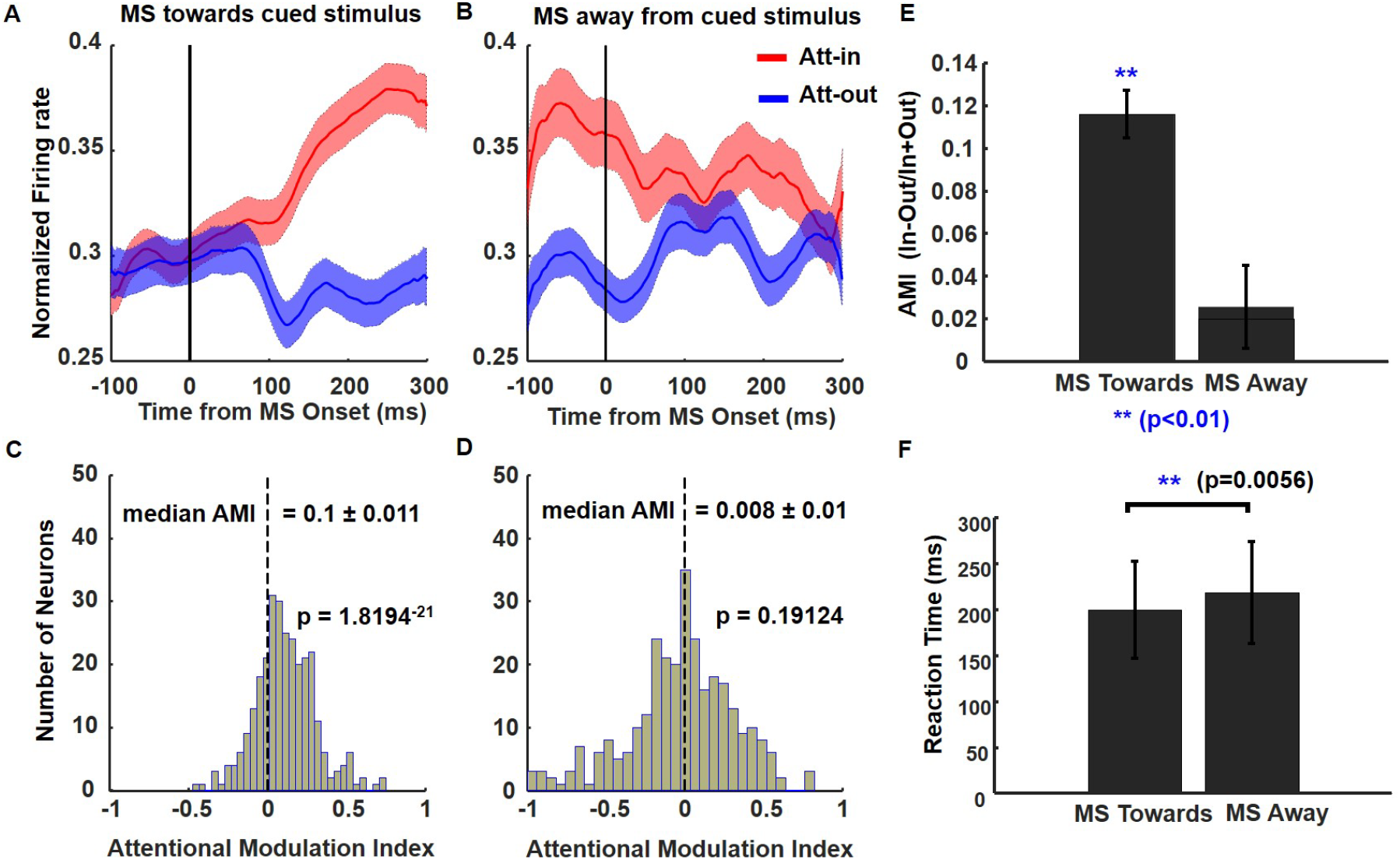
V4 Population activity for the last MS triggered interval preceding target color change and behavioral performance. (A and B) Same analysis as in Figure 2 except that the normalized firing rates are locked to the last MS detected before target color change happens. We observe that there is still significant attentional enhancement for the MS towards the target in the RF. (C and D) Same analysis of AMI but only for the last MS. (E) Summary of the AMI for both MS towards and MS away for the last MS. There is significant attentional enhancement for MS towards the RF even for the last MS before change detection. Reaction times to saccade to the target following target color change with respect to the last MS being towards or Away. The reaction time to saccade to the target was on average 20ms faster if the last MS was towards the target than away. *p=0*.*0056* was calculated via two sample Kolmogorov-Smirnov test. See Supplementary Figure S6 for histograms of reaction time distribution and the cumulative distribution of their response latency.

As in V4, population activity of neurons recorded in the lateral nucleus of Pulvinar (Figure S5 E-H) also showed enhanced attentional modulations on the first MS for the MS towards direction (*t-test, n=339, p=4*.*92*^*-4*^, *median AMI = 0*.*02±0*.*007)*, and not to MS away (*t-test, n=339, p=8*.*01*^*-9*^, *median AMI = -0*.*04±0*.*01)*. Unlike in V4, even in the subsequent MS intervals (excluding the first MS) the attention effects were only significant in the MS-towards condition (Figure S5 I-L). However, the attentional effects were weaker and noisier than in V4 which makes it difficult to draw strong conclusions. Despite this, there was a significant positive AMI for MS towards the cued stimulus in both the first MS and subsequent MS intervals (Figure S5 M-N). The overall effects of enhanced attentional modulation for all MS intervals are presented in Figure S5 A-D.

A similar analysis of the population activity of neurons recorded in area IT (Figure S6 E-H) also showed enhanced attentional modulations for the first MS directed towards (*t-test, n=357, p=1*.*74*^*-08*^, *median AMI = 0*.*044±0*.*01)*, and not for the first MS directed away (*t-test, n=357, p=9*.*72*^*-13*^, *median AMI = -0*.*085±0*.*01)*. Enhanced attentional modulation persisted for the subsequent MSs excluding the first MS as well when the MS was directed towards the target (Figure S6 I-L). There was significant enhancement for MS towards the target for both the first MS and when the first MS was excluded (Figure S6 M-N) as well despite the fact that the RF of the IT cells encompassing the entirety of the stimulus array in the contralateral visual field.

### Object Decoding of V4 neurons during full stimulus period versus different MS onset intervals reveals distinct regimes

In (Lowet et al., 2018) we used a simple analysis of variance (ANOVA) of the neuronal firing rates of V4 neurons as a function of attention and MS onsets at the RF location, which revealed that the ability to discriminate the objects was significantly greater when the MS was directed towards the attended stimulus. Here, we extend this analysis to look at the temporal profile of the performance of a linear decoder trained on the population of V4 neurons for both the full stimulus periods and the MS triggered intervals.

We first considered the time-course of the effects of attention on decoder performance over the full stimulus interval, but without taking MSs into account. We first trained a linear decoder from the activity of individual V4 neurons from the cue-first/stim-second sessions (*n=132*), for the seven distinct objects shown in the stimulus array in a 1000ms time window beginning at stimulus onset. For the cue-first, stimulus second condition, the transient sensory response occurred after the attentional cue, so the transient response was included in the data entered into the decoder. If the decoder were to perform at chance levels for discriminating the seven objects, its performance would be (100/7=14.3%) irrespective of the time intervals chosen.

The average temporal profile of the decoder performance, after a ten-fold cross-validation (*80% training + validation set, the remaining 20% was held-out for testing*) for the transient sensory response, not taking MS into account, is shown in Figure 7A. The decoder’s performance for object categories during the transient response were both significantly above chance levels for both Att-in and Att-out conditions. The ability to discriminate object categories for the Att-in condition reached a maximum of ∼60% at around 200ms after stimulus onset and fell to near chance by around 500 ms after stimulus onset. The decoder performance for Att-out trials was also significantly above chance, reaching a maximum of ∼43% to discriminate the seven objects. This result establishes that if the time period just after stimulus onset is considered, there is enough information in the firing rates of V4 neurons to discriminate objects, and the discrimination is consistently better if attention is directed towards the RF than outside. This result is consistent with decoding results in IT cortex shown in (Zhang et al., 2011).

**Figure 7.**
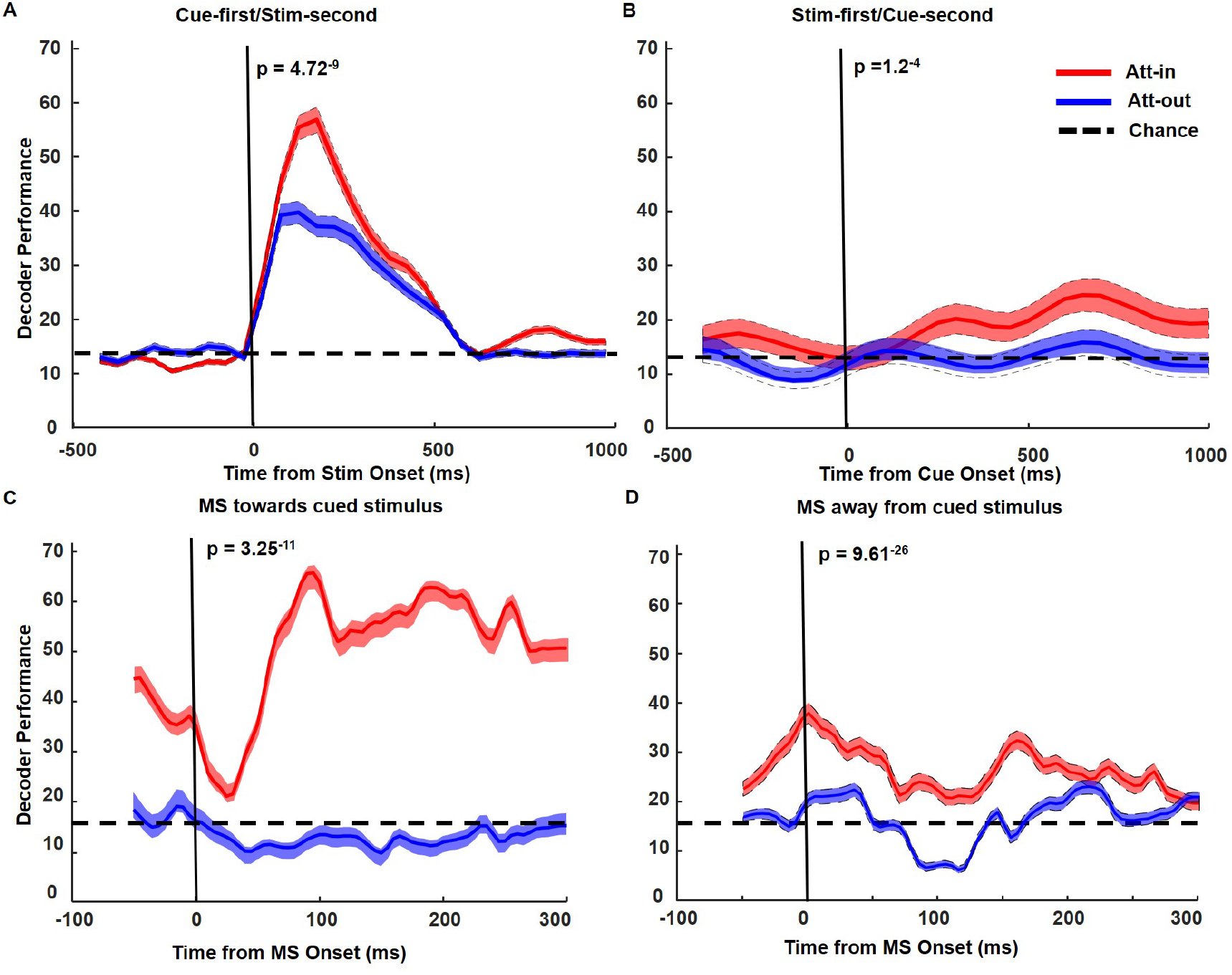
Object decoding of V4 neurons during full stimulus time period and MS triggered intervals. A) Linear decoder performance for object categorization by V4 neurons (*n= 132*) from the cue first/stimulus second sessions including the transient stimulus onset period in the 1000ms following stimulus onset. B) Linear decoder performance for object categorization by V4 neurons (*n= 150*) for 1000ms during the stimulus first/cue second condition, which does not include a stimulus transient. C) Linear decoder performance for the 300ms time periods for both Att-in and Att-out immediately after an MS is directed towards the cued stimulus (*n=282*). Decoder performance was significant for MS towards intervals directed to the RFs of V4 (All the MS post stimulus and spatial cue presented were included). D) Same as in (C), but when the MS is directed away from the cued stimulus.

If we excluded the initial transient response after stimulus onset, the decoder performance diminished substantially. To calculate the decoder performance without the stimulus transient, we used the data from the stimulus first-cue second condition (*n=150*), and we used the data for the 1000ms interval following the onset of the attentional cue, which was outside the RF. In this case, the transient response to the stimulus was over by the time the cue is presented. The temporal profile of the decoder performance trained for this interval are shown in Figure 7B. Unlike in Figure 7A, which included the transient stimulus, the decoder’s performance for object categories was at chance levels for Att-out conditions, i.e., when the cued target was out of the V4 RFs, the ability of the V4 neurons to discriminate objects within the RF was at chance throughout the trial. By comparison, the decoder performance for Att-in was significantly above chance from around 100-200 ms after the cue. While significantly above the Att-out condition, and chance (14.3%), the decoder performance for Att-in without taking MS into account was only marginal, reaching a maximum of ∼23% to discriminate the seven objects. This is in contrast to the nearly ∼60% for Att-in to decode when the transient response was included (see Figure 7A). Thus, when excluding the stimulus transient, and only looking at the stable/attention period, we observe that there is only a marginal boost to object decoding with attention directed to V4 RFs.

We then computed the decoder performance when taking MSs into account. We analyzed the intervals following the four different attention and MS direction conditions, for the 300ms time window after the MS onset. Because the number of trials and stimuli needed for the decoding analysis were not sufficient for the first MS alone, we conducted the analysis on all of the MS intervals, and then separately conducted the analysis with the first MS interval excluded (Figure S8). We first present the results for all MS intervals.

The time average of the decoder performance after cross-validation for attention periods when the MS was cued towards the stimuli and away from the stimuli are shown in Figures 7C and 7D, respectively. The decoder performance was significantly better for the attention-in condition following the MS towards the cued target in the RF (Figure 7C) in comparison to when it was cued to the target away from the RF (which remained below chance), and remained so for the entire 300ms time window (*t-test, n=282, p=3*.*25*^*-11*^). Similarly, the decoder performance for the MS away from the cued target in the RF (Figure 7D), while lower than for the MS towards the cued target in the RF, was still significant in comparison to when the MS was directed away from the target outside the RF (which remained below chance as well), and remained so for the entire 300ms time window (*t-test, n=282, p=9*.*61*^*-26*^). The overall temporal profile for performance for Att-in MS towards had a peak at ∼67% around 100ms after MS onset, meanwhile the maximum decoder performance for Att-in MS away peaked at ∼35%. Thus, it seems as though the Attention-in MS towards condition caused a transient “burst” of improved decoding performance, reminiscent of the decoder performance following the stimulus transient, while for all other conditions the performance was near chance or just above chance (Attention-in MS away condition).

This interpretation was supported by comparing the mean classification accuracy for the transient stimulus period (Figure 7A), the full stimulus attention period (Figure 7B), and the MS triggered intervals (Figure 7C-D) for held-out/test data (20% of the dataset) presented in Figure 8. Att-in MS towards had the highest classification accuracy on the held-out test data (∼51%), followed by the transient stimulus presentation period (with Att-in = ∼45%; Att-out =∼36%), and Att-in MS away (∼28%). The classification accuracy for the Att-out MS intervals were only marginally above chance (∼15%, and ∼17% respectively). This decoding performance was further tested using an ANOVA computed on the mean firing rate of the population in time windows locked to the transient stimulus period (400ms) after cue appeared, the full stimulus stable/attention period (1000ms), and the MS triggered intervals (300ms). The results showed a significant effect of attention (as quantified by the F-value being much greater than 1) for decoding in the transient stimulus period after cue presentation (Att-in: *t-test, n=132, F-value= 4*.*1;* Att-out: *t-test, n=132, F-value= 2*.*3*), for Att-in during the MS triggered intervals (MS Towards: *t-test, n=282, F-value= 8*.*8*; MS Away: *t-test, n=282, F-value= 1*.*8*); a non-significant effect for the full stimulus stable/attention period (Att-in: *t-test, n=282, F-value=0*.*33;* Att-out: *t-test, n=282, F-value=0*.*45*), and for the Att-out trials during the MS triggered intervals (MS Towards: *t-test, n=282, F-value= 0*.*95*; MS Away: *t-test, n=282, F-value= 1*.*1*).

**Figure 8.**
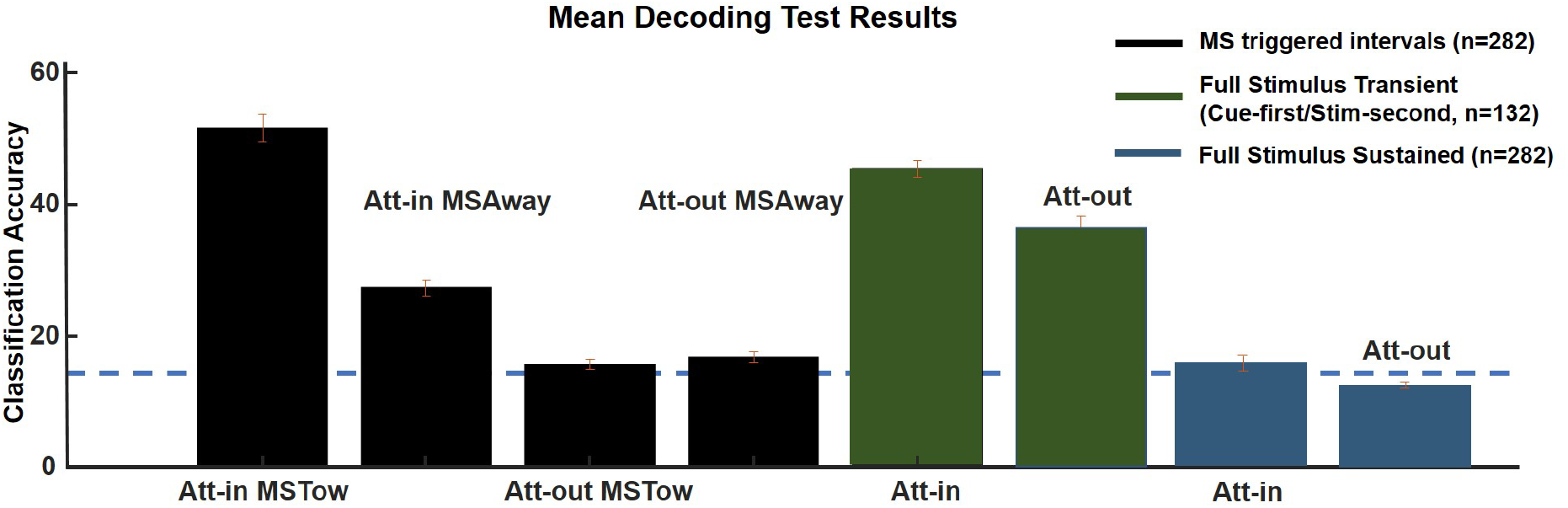
Comparison of classification accuracy across conditions.

Similar results were found when we excluded the first MS (Figure S8 A-C). When the MS was directed towards the cued stimulus, the decoder performance for the attended stimulus rose over a period from 50-100ms after the MS, and then remained constant for the remainder of the interval, whereas the decoder performance for the unattended stimulus remained at about chance. Correspondingly, the mean decoder performance for the attended stimulus (peak at ∼ 69%) was significantly greater than chance and significantly greater than for the unattended stimulus (peak at ∼17%) (*t-test, n=282, p=1*.*89*^*-14*^) (Figure S8 A). By comparison, in intervals with the MS directed away from the RF, the decoder performance for the attended stimulus was above chance (peak at ∼ 37%) and was significantly greater than for the unattended condition (peak at ∼22%) which was near chance levels (*t-test, n=282, p=2*.*32*^*-7*^) (Figure S8 B)

The classification accuracy for held-out/test data was similar to that of all MS trials (Figure S8 C, compare with Figure 8). Since these are linear decoders, and the decoder performance for all MS is the average of first MS and subsequent MS, it seems likely that the decoder performance after the first MS only would be similar to that of the all MS condition. Thus, after the first MS, the effects of attention on firing rates are somewhat maintained for subsequent MS in either direction, but the effects of attention and MS direction on decoding performance continues to be pronounced.

What was responsible for the better decoding performance with attention in the MS towards and away from the RF conditions? One possibility is that cells were more selective for different stimuli during these conditions, i.e. cells had sharper tuning curves. To test this idea, we examined the object tuning curves under the different conditions. We first computed the normalized mean of the firing rate activity for the transient stimulus period after cue (50-350ms after stimulus onset), the full-stimulus period without transient (1000ms time window), or the MS triggered intervals (300ms time window) for each of the 7 objects within its RF, and then rank ordering them from most preferred to the least preferred object. The greater, and higher the slope of the curve, the greater the selectivity. In Figure 9, the relative firing rates to the most to lease preferred stimuli in the different conditions are shown for the V4 population.

**Figure 9.**
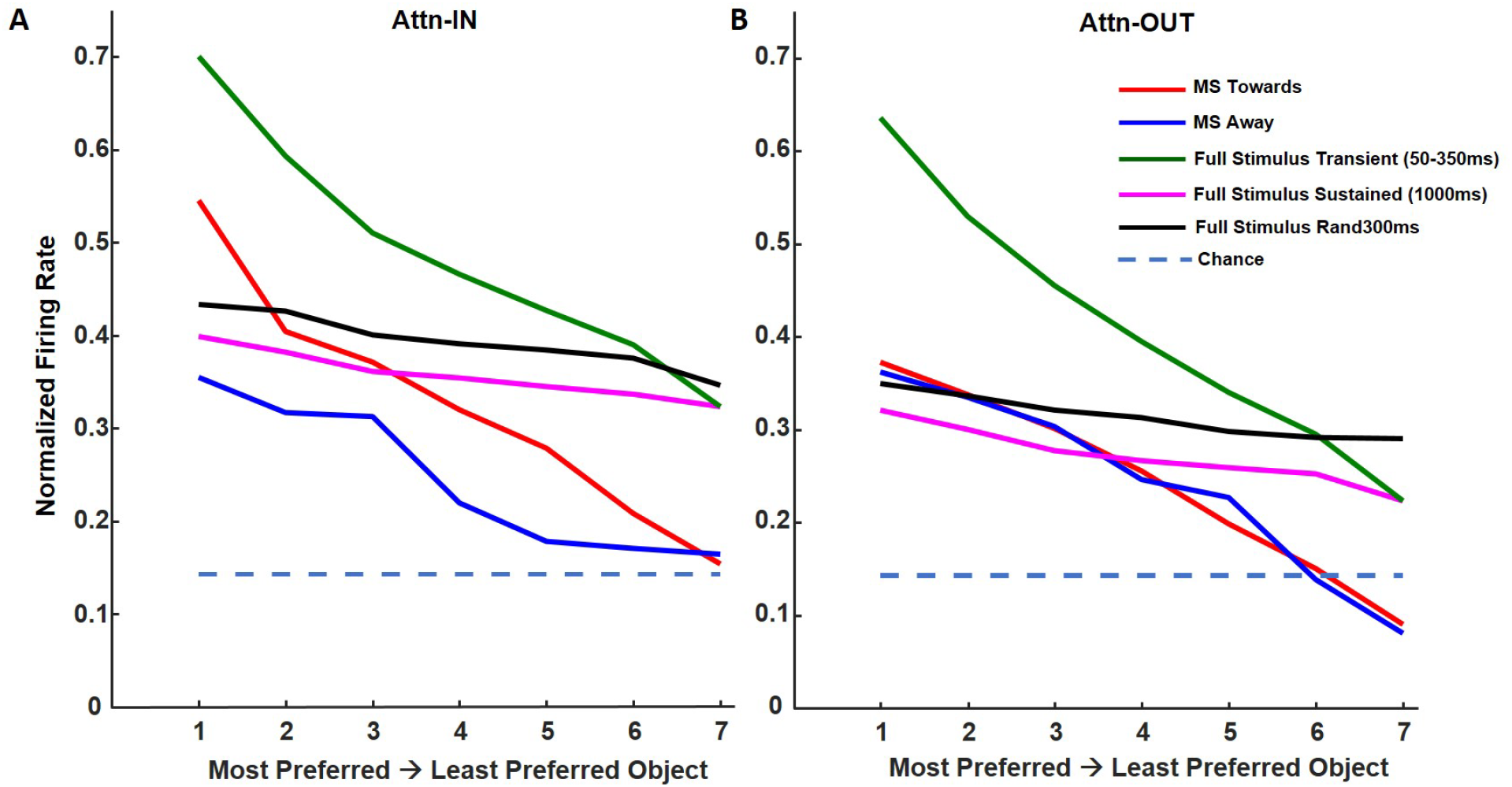
Object Selectivity of V4 neurons across the population for full stimulus vs MS triggered intervals (All MS were included). A) Mean normalized population firing rates of V4 neurons *(n=282)* to the Att-in condition ranked from the most preferred to the least preferred objects reflecting different object tuning profile for the neurons for both full stimulus and MS triggered intervals. B) Same as in (A) but for the Att-out condition. See Supplementary Figure S8 (D-E) for the same analysis done but excluding the first MS. See Supplementary Figure S9 for example of differential object selectivity in exemplar V4 neurons depending on MS type.

Considering first the attention-in tuning curves (Figure 9A), the turning curves for the Full Stimulus Transient (not taking MS into account) and MS triggered had the steepest slopes, consistent with the decoding results. The flattest slope was for the Full-Stim conditions excluding the transient response. The overall magnitude of response to the preferred stimulus was largest for the Full-Stim Transient response and the MS-Towards conditions, with a suggestion that the biggest increase in response occurred to the most preferred stimulus following the MS-towards. Thus, the differences found among conditions in the decoding results are overall consistent with the differences in tuning curves. They suggest that, with attention, sensory coding is optimal for the initial transient response to the stimulus, and it becomes substantially weaker for the more sustained phase of the response. MSs restore sensory coding performance during the sustained phase of the response, with the biggest effect for the MS-towards the attended stimulus.

Considering next the attention-out tuning curves, (Figure 9B), the tuning curve for the Full Stim Transient condition had the steepest curve and highest firing rate, similar to the slope of the tuning curve for the attend-in condition but with somewhat smaller responses. The similarity in tuning curves across attention conditions is consistent with prior studies showing that spatial attention does not sharpen tuning curves in V4, based on average firing rates (David et al., 2008; Luck et al., 1997; McAdams and Maunsell, 1999, 1999; Ogawa and Komatsu, 2004; Reynolds et al., 2000; Williford and Maunsell, 2006). The slopes of the tuning curves for both the MS-towards and MS-away tuning curves were similar to the curve for the stimulus transient response, but the overall responses were much smaller. It appears that in the attention-out condition, a MS in either direction helps to restore tuning. The flattest tuning curves were for Full Stimulus conditions excluding the transient. The corresponding scatter plots for the Z-scored mean firing rates (during the 300ms) for all the neurons in order of object preference for MS-towards, MS-away (for all MS intervals) and full stimulus are presented in Figures S7. Similar results were found for responses excluding the first MS (Figure S8 D-E), similar to what we found with the decoding results. The scatter plots for the Z-scored mean firing rates for the MS intervals excluding the first MS towards or away are presented in Figure S8 F and G respectively.

In sum, the tuning curves for the transient response following stimulus onset are among the sharpest compared to all conditions, and the firing rates overall are higher. The effects of attention are modest for the transient response. Following the transient response, firing rates are reduced and tuning curves flatten out when MSs are not taken into account, consistent with the finding that stimulus decoding is poor following the stimulus transient. The interval following a MS seems to restore the sharpness of the tuning curve, and the influence of attention seems to substantially increase the firing rates for the preferred stimulus as well, all consistent with the good decoding performance, especially in the MS-Towards, Attention-in condition. Attention and MSs towards the attended stimulus thus counteract the reduction in tuning found during the sustained phase of the response.

Thus, the direction of MS during attention leads to distinct object selectivity regimes for every MS interval, modulating both stimulus representations and tuning. This result stands in contrast to what we found for the effects of attention on average firing rates, namely that attention enhanced firing rates for both MS directions after the first MS. This differential object selectivity depending on the MS types is shown for a few V4 single neurons examples in Figure S9. Such changes in object selectivity in V4 might be useful for downstream areas like IT, resulting in better object categorization. In Lowet et al., (2018) we had already shown evidence that IT firing rates increase after MS towards than for MS away.

## DISCUSSION

The results show that MSs significantly affect the neural processing of object features with attention. We found the largest effects of MS direction on firing rates during the first MS after stimulus or cue onset. On that first MS, attending to the stimulus in the RF was strongly modulated by MS direction, with little or no effect of attention when the MS was directed away from the attended stimulus. On subsequent MSs, attention enhanced the response to the RF stimulus during both MS directions, but the largest effects were still for MS directed towards the attended stimulus. Comparable results were found in the SC, where the effects of attention are strongest during MSs directed towards the attended stimulus but there remains a significant effect of attention even with MS directed away from the attended stimulus (Yu et al., 2021). It was suggested based on the SC results that MS direction is a marker for the locus of attention, but the MS is not the cause of attentional modulation in the cortex. We favor a somewhat different interpretation, which is that spatial attention and MSs may have independent effects on neuronal responses. We found that attentional deployment to the cued stimulus is delayed until the first MS towards that stimulus, consistent with the findings of (Lowet et al., 2018). Here, we found that if the first MS was directed away from the cued stimulus in V4, the effects of attention were delayed until the second MS, which was towards the cued stimulus. Once attention is deployed, it may persist at the site of the cued stimulus to a certain extent, even when a later MS is directed back to the fixation target, which would explain why we found a certain degree of attentional enhancement on all MSs subsequent to the first MS. If a MS directed towards the cued stimulus means that there is more attention directed to the cued stimulus compared to when the MS is directed away, then one could say that the direction of the MS is a marker of attention. On the other hand, if attending to a stimulus and making a MS towards a cued stimulus have independent effects on neuronal responses, then the MS direction could be both a marker and a cause of the attentional effects. Our findings on the effects of MS on stimulus encoding support this latter interpretation.

It is possible that the onset of the first MS towards the cued stimulus might trigger a global and critical change in the dynamic regime enabling attentional modulation across these brain areas (Hesse and Gross, 2014; Sergent et al., 2021). In the language of dynamical systems theory, the first MS towards a target represents a bifurcation that qualitatively pushes the neural dynamics from one state (no-attention) to another (attention) (Cocchi et al., 2017; Müller et al., 2020; Scheffer et al., 2012; Toker et al., 2022). However, with the subsequent MSs, the attentional firing rate enhancement perseveres.

In contrast to the differential effects on firing rates on the first and subsequent MSs, both the first MS and subsequent MSs profoundly modulate stimulus encoding and stimulus representations, presumably accounting for the enhanced object representations found in downstream areas such as IT cortex (Lowet et al., 2018). In line with enhanced object processing, we found that MSs also improved the reaction time to detect color changes of the cued stimulus. Previous studies have reported only multiplicative effects with no sharpening of V4 tuning curves during spatial attention (McAdams and Maunsell, 1999; Luck et al., 1997; Reynolds et al., 2000) while response shift (tuning shift) has been found during feature attention (Bichot et al., 2019; Ipata et al., 2012; Maunsell and Treue, 2006; Nobre et al., 2006; Zhou and Desimone, 2011). We found here that during intervals triggered by a MS directed towards the attended stimulus, there is both response gains and response shifts (sharpening of tuning curves) for object/feature processing (Motter, 1994, 1993).

A possible mechanism for these response shifts is suggested by our finding that stimulus selectivity is much sharper during the initial transient response to the stimulus onset (included in the cue-first, stimulus-second condition) compared to the longer sustained response (included in the stimulus-first, cue-second condition). MSs may “refresh” the stimulus representation during prolonged fixations, as has been suggested by others (Martinez-Conde et al., 2004; Martinez-Conde et al., 2009; Martinez-Conde et al., 2013; Engbert, 2006). MSs in either direction improve stimulus decoding compared to the sustained response, but we found the biggest effect occurred when the MS was directed towards the attended stimulus, suggesting that the “refreshing” created by the MS interacts with the effects of attention in some multiplicative form.

Several theories have suggested that covert spatial attention is a rhythmic sampling process associated with a 3-4Hz theta rhythm of the local field potentials (Bosman et al., 2009; Fiebelkorn et al., 2013; Fiebelkorn and Kastner, 2020; Helfrich et al., 2018; Schroeder et al., 2010; Schroeder and Lakatos, 2009; Song et al., 2014; Spyropoulos et al., 2018). Here too, we observe attentional modulation is either enhanced or reduced depending on the direction of the MS. MSs have been shown to powerfully modulate neural firing rates and LFPs across the visual system (Leopold and Logothetis, 1998; Martinez-Conde et al., 2013, 2009), tightly linked to covert attentional modulations (Chen et al., 2015; Denison et al., 2019; Hafed, 2013) and therefore a plausible mechanism for regulating brain-wide state transitions during covert attention.

We continue to use the word “covert” to study spatial attention, as if it is entirely separable from the oculomotor systems. We believe this is a misnomer. Our work adds to a growing set of studies showing that separation of the physiological mechanisms that mediate eye movements from those that mediate the effects of attention seriously misrepresents the intricate and dynamic interplay between the two systems. After all, visual processing is necessary for action and eye movements provide the most important form of selecting behaviorally relevant targets to visually process. While most studies have looked at how saccades help with such a visual selection mechanism via shifts in attention, MSs are usually considered to be subliminal movements typically, if not treated outrightly as involuntary. This too deserves reconsideration. Zuber and colleagues showed that in terms of their velocity, amplitude, and direction distributions, MS and saccades share a common profile and likely share common physiological mechanisms (Zuber, et al., 1965). The evidence that MSs are generated by saccade generating areas is now well established (Bollimunta et al., 2018; Hafed and Krauzlis, 2012; Peel et al., 2016). If so, is the relationship between MS and covert spatial attention similar to saccades and feature attention, in that they are normally coordinated by the same circuit, but can be dissociated under some conditions (Bichot et al., 2005; Kowler et al., 1995)? Further experiments along these lines will help us establish the dynamic relationship between the attentional systems and the oculomotor systems towards synthesizing an active sensing and attentional sampling (Schroeder et al., 2010; Schroeder and Lakatos, 2009; VanRullen, 2016, 2013) view of cognition.

## ACKNOWLEDGMENTS

Supported by NIH R01-EY029666.

## Supplementary Figures

**Figure S1.**
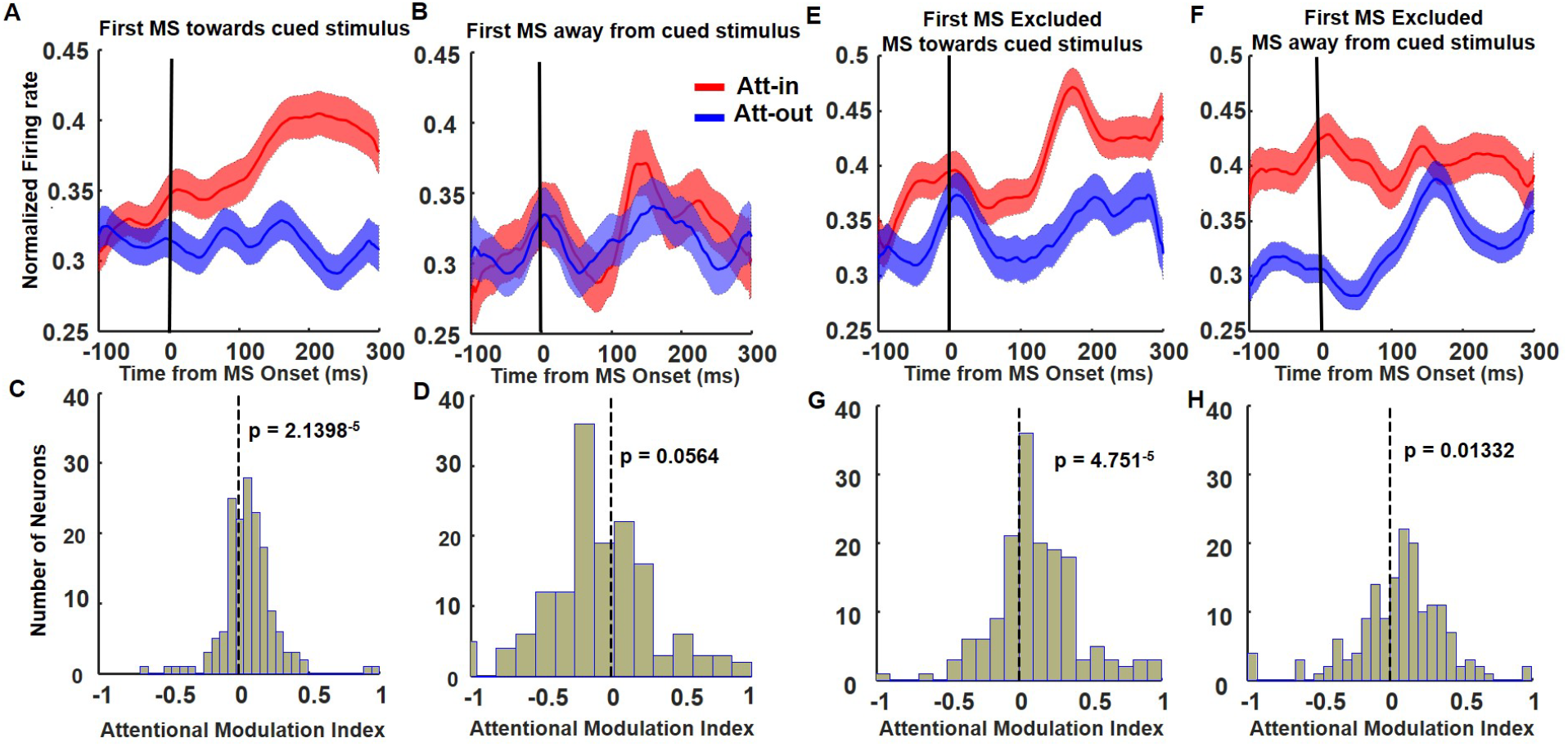
The role of the first MS, and subsequent MSs during the attention period in area V4, but only for the stim-first/cue-second trials *(n=161 neurons)* combined across monkeys and sessions (*k=18*) (A-D) Schematic same as in Figure 2 for the first MS during attention period. (E-H) Schematic same as in Figure 4 excluding the first MS.

**Figure S2.**
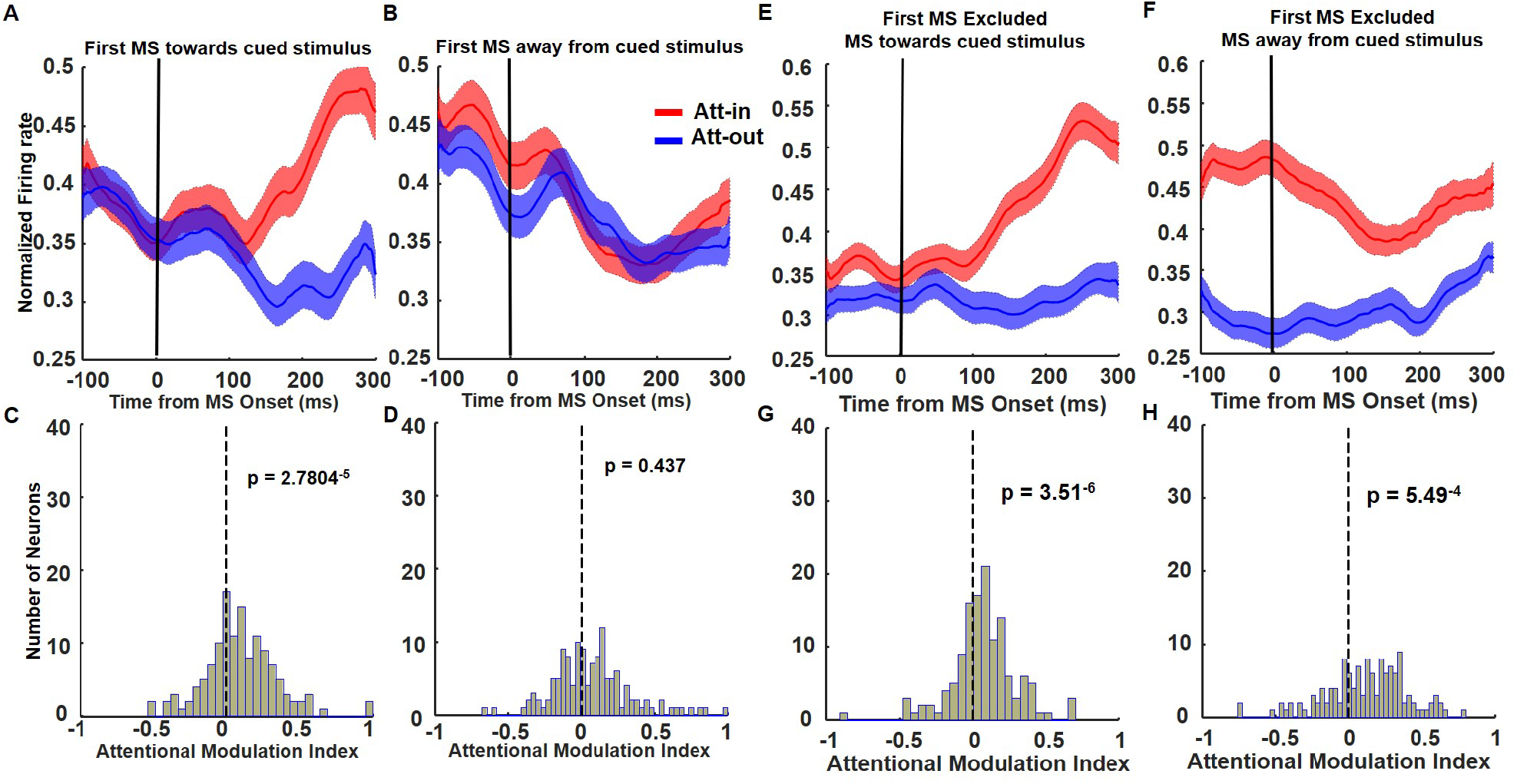
The role of the first MS, and subsequent MSs during the attention period in area V4, but only for the cue-first/stim-second trials *(n=132 neurons)* combined across monkeys and sessions (*k=16*) (A-D) Schematic same as in Figure 2 for the first MS during attention period. (E-H) Schematic same as in Figure 4 excluding the first MS.

**Figure S3.**
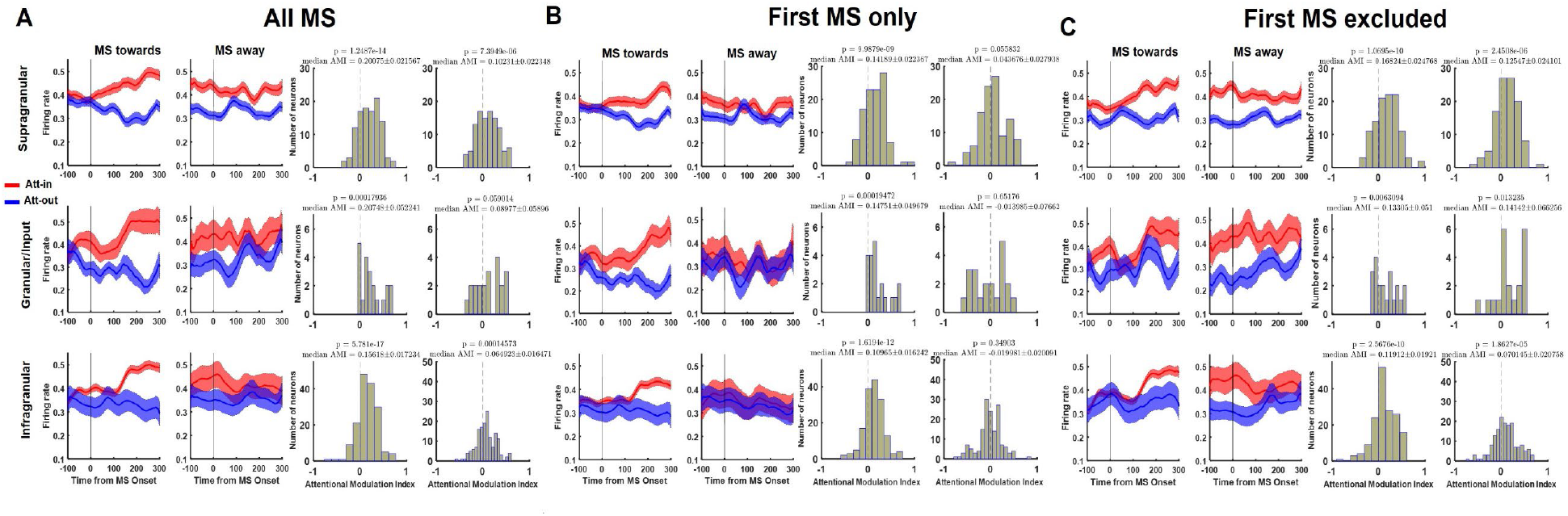
Laminar profile of MS triggered V4 firing rates during the attention period. (Top row) Normalized firing rate during MS onset intervals (towards and away), and the corresponding AMI histogram distributions for the cells in the supragranular layer (*n= 109*) for all MS intervals (A), first MS only (B), and first MS excluded (C) respectively. The median AMI values and whether they are significant between Att-in and Att-out are provided for each condition at the top of the AMI histograms. (Middle row) The same convention of (A-C) as above, but only for the granular/input layer (*n=21*). (Bottom row) The firing rates, and AMI histogram distributions for cells in the infragranular/deep layers (*n=163*). See Figure 5 for overall summary of MS related attention effects across the laminar profile of V4.

**Figure S4.**
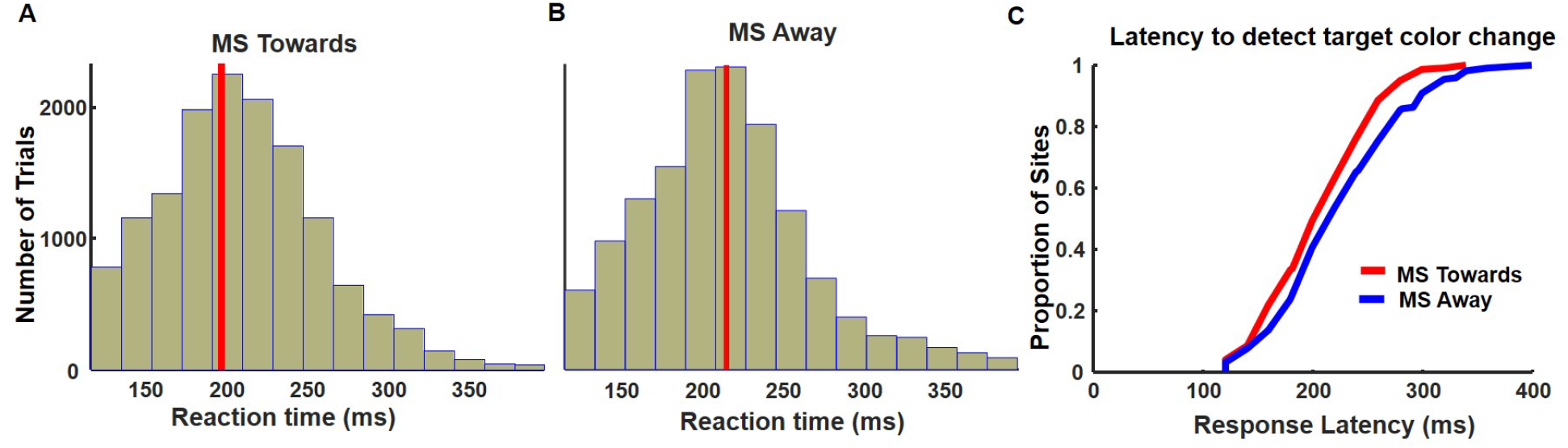
Reaction time (RT) distribution to saccade to the target as a function of the last MS before target color change. A) RT distribution to saccade to the target if the last MS was towards the target before color change. B) RT distribution if last MS was away from the target. The red vertical lines in both figures marks the mean RT value (in ms). C) The empirical cumulative distribution function (ecdf) for response latency for the last MS towards vs MS away before color change for the target in the V4 RF. See Figure 6 for V4 firing rate analysis and RT summaries.

**Figure S5.**
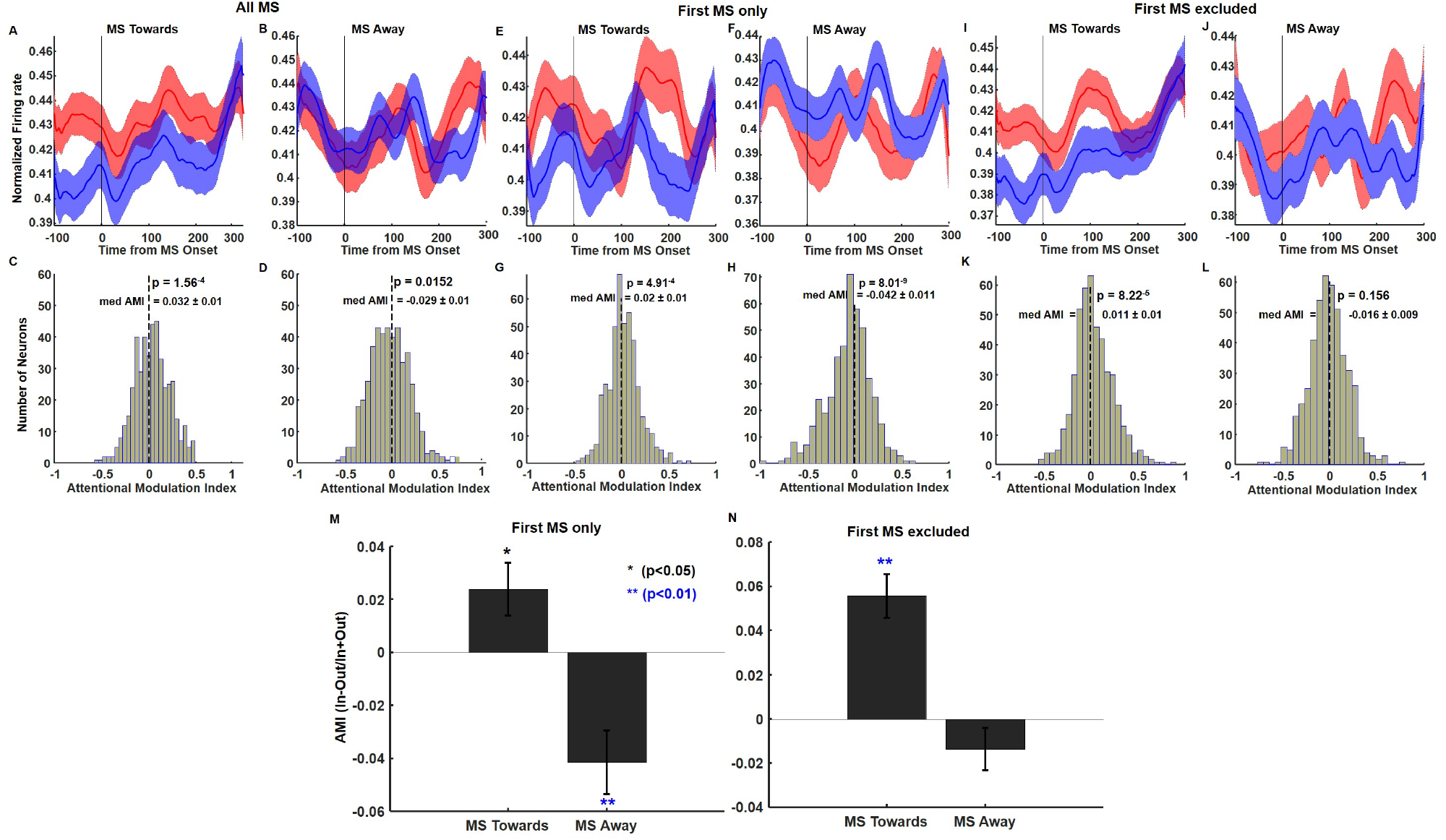
Population firing rate of pulvinar neurons during MS triggered intervals (*n =339*). The schematic is similar to the previous figures for MS towards and MS away during attention. The median AMI values and whether they are significant between Att-in and Att-out are provided for each condition at the top of the AMI histograms. (A-D) Normalized population firing rates and AMIs for all MS intervals post stimulus and cue onset within and outside the RF. (E-H) Same as (A-D) but only for the first MS (compare with Figure 2 and S3B for V4). (I-L) Firing rates and AMIs when the firs MS is excluded (compare with Figure 4 and S3C for V4). (M) Summary of the population AMI response of pulvinar neurons to only the first MS. (N) Summary of the population AMI with the first MS excluded (compare M and N with Figure 5).

**Figure S6.**
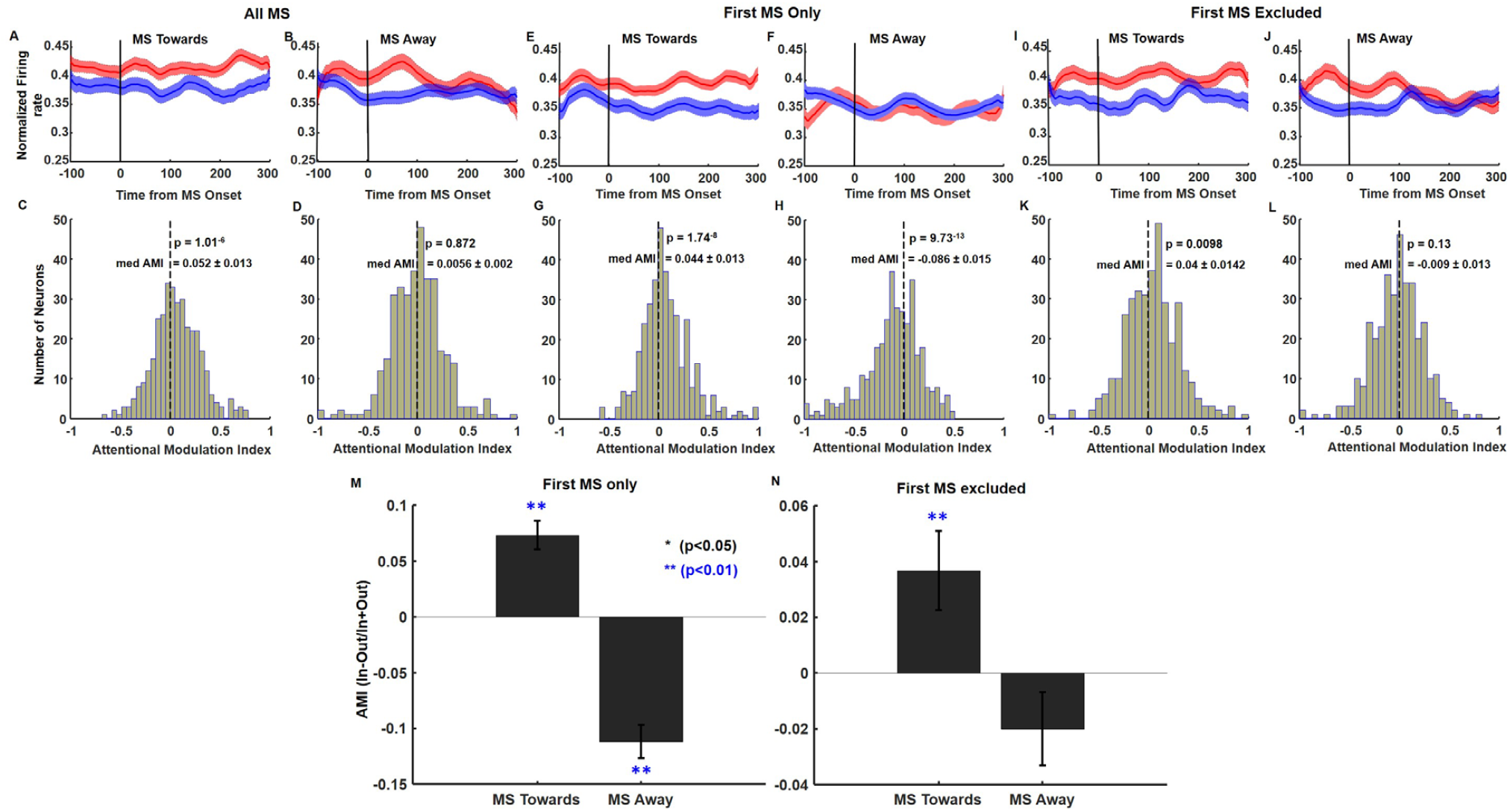
Population firing rate of neurons in area IT during MS triggered intervals (*n =357*). Schematic same as in Figure S5. (A-D) Normalized population firing rates and AMIs for all MS intervals during Attention within and outside the RF. (E-H) Same as (A-D) but only for the first MS. (I-L) Firing rates and AMIs when the firs MS is excluded. (M and N) Summary of the population AMI response to only the first MS, and the first MS excluded respectively.

**Figure S7.**
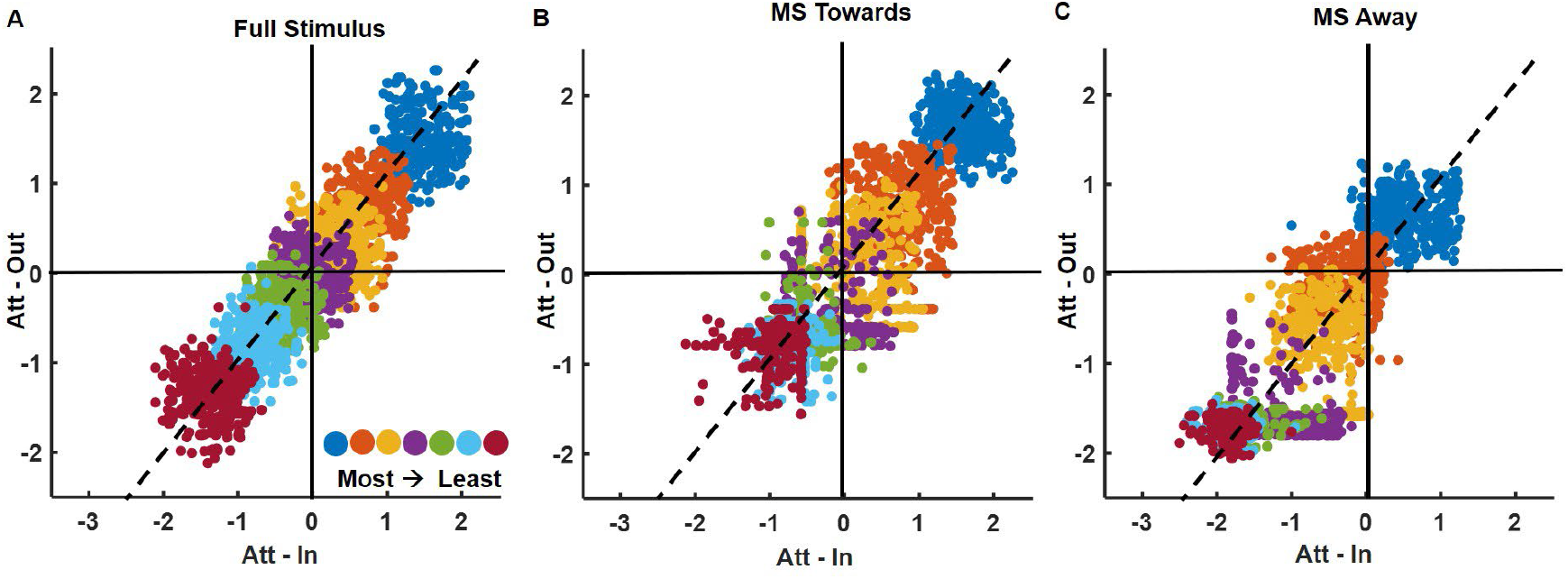
Scatter plots of the Z-scored mean firing rates for all V4 neurons (*n=282*) in order of object preference. A) For the full stimulus attention period. B) For all MS towards the target intervals (300ms). C) For all MS away from the target (see Figure S8 F-G when the first MS is excluded).

**Figure S8.**
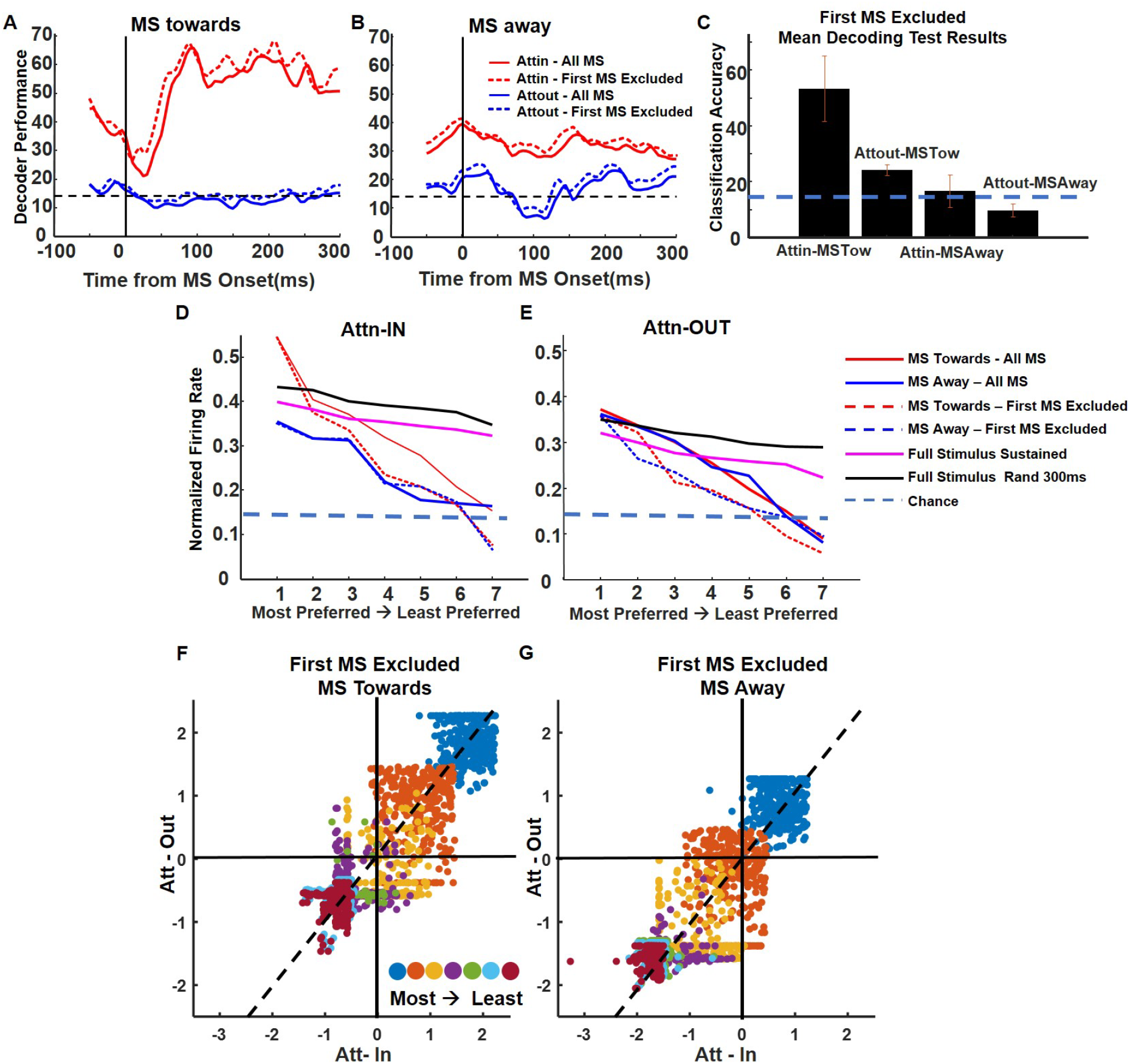
Object decoding with V4 neurons (*n=282*) excluding the first MS post stimulus and cue presentation. A) Linear decoder performance for the 300ms time periods for both Att-in and Att-out immediately after an MS is directed towards the cued stimulus. Decoder performance was significant for MS towards intervals directed to the RF. B) Same as in (A), but when the MS is directed away from the cued stimulus. There was still a significant decoder performance for MS away from the RF, but lower than when the MS was directed towards. (compare with Figure 7C and D). C) Mean decoding results on the held-out test data with the first MS excluded showed similar classification accuracy as when all MS intervals were used for decoding (compare with Figure 8). D) Mean normalized population firing rates to the Att- in condition ranked from the most preferred to the least preferred objects reflecting object selectivity and tuning during MS triggered intervals. E) Same as in (D) but for the Att-out condition (compare with Figure 9). F) Scatter plots of Z-scored firing rates in order of object preference in MS towards intervals. G) Same as in (F) but for MS away intervals (compare with Figure S7 B and C).

**Figure S9.**
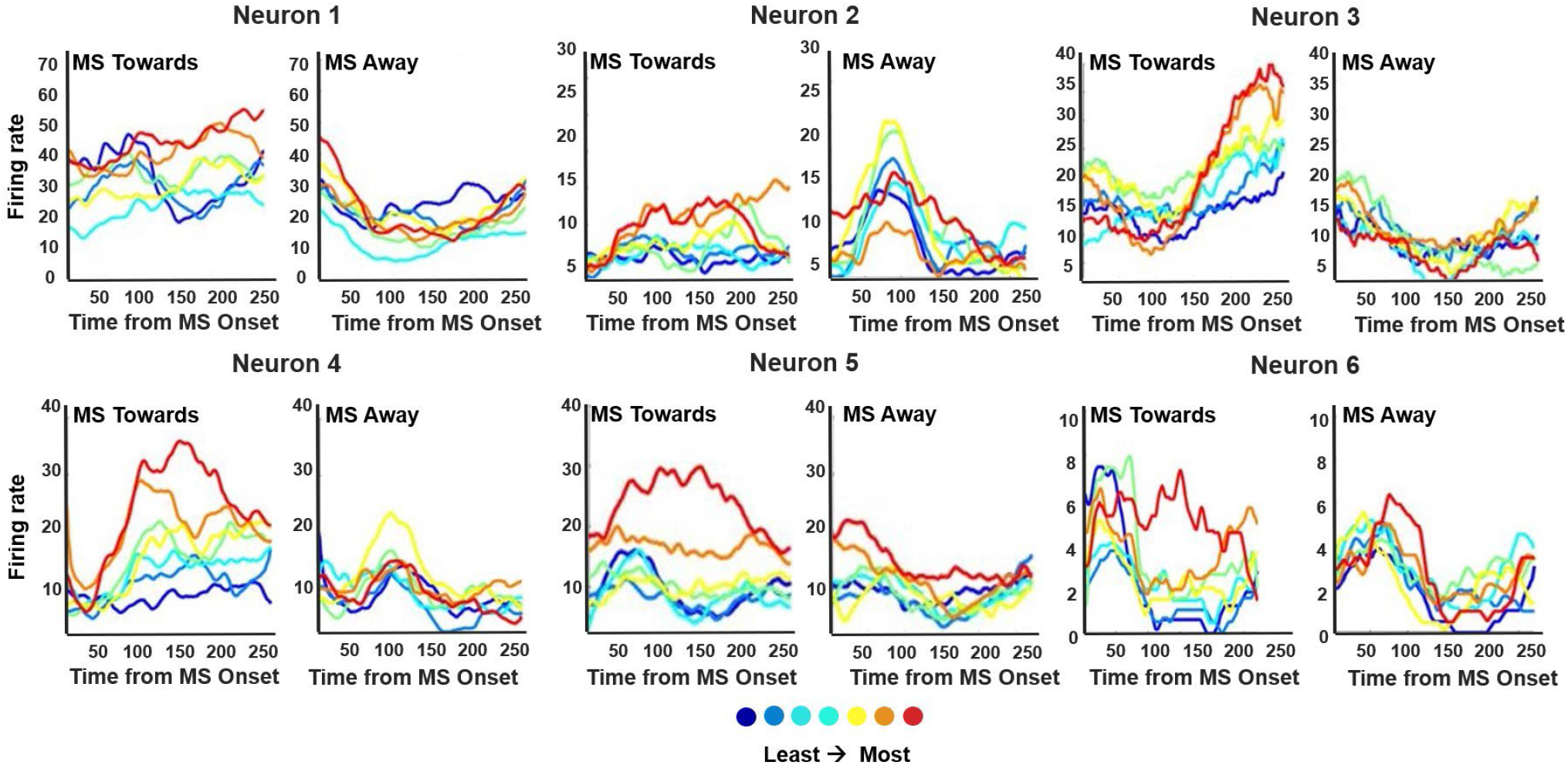
Differential object selectivity/preference depending on the MS type for Attention-In is shown for a few example neurons in V4.

